# Autophagy determines osimertinib resistance through regulation of stem cell-like properties in EGFR-mutant lung cancer

**DOI:** 10.1101/330092

**Authors:** Li Li, Yubo Wang, Lin Jiao, Caiyu Lin, Conghua Lu, Kejun Zhang, Chen Hu, Junyi Ye, Dadong Zhang, Mingxia Feng, Yong He

## Abstract

Drug resistance to Osimertinib, a 3^rd^-generation EGFR-TKI is inevitable. Autophagy plays a contradictory role in resistance of 1^st^ and 2^nd^ generation EGFR-TKI, and its significance in osimertinib resistance is much less clear. We therefore investigated whether autophagy determines osimertinib resistance. First, osimertinib induced autophagy to a much greater extent than that of gefitinib, and autophagy inhibition further increased osimertinib efficacy. Next, enhanced autophagy was found in osimertinib resistant cells and autophagy inhibition partially reversed osimertinib resistance. Enhanced stem-cell like properties were found in resistant cells, and siRNA-knock down of *SOX2* or *ALDH1A1*reversed osimertinib resistance. Of note, autophagy inhibition or siRNA-knock down of Beclin-1 decreased expression of SOX2 and ALDH1A1 and stem-cell like properties. Next, autophagy inhibition and osimertinib in combination effectively blocked tumor growth in xenografts, which was associated with decreased autophagy and stem cell-like properties *in vivo*. Finally, enhanced autophagy was found in lung cancer patients with resistance to osimertinib. In conclusion, the current study delineates a previously unknown function of autophagy in determining osimertinib resistance through promoting stem-cell like properties.

## Introduction

Non-small-cell lung cancer (NSCLC) treatment has evolved dramatically in the last decade, from the traditional “one-size-fits-all” chemotherapeutic approach to new targeted therapies against oncogenic driver mutations. In NSCLC patients with EGFR-activating mutations, 1^st^-generation epidermal growth factor receptor tyrosine kinase inhibitors (EGFR-TKIs) have become the standard first-line therapy with dramatic therapeutic efficacy (Nguyen & Neal, 2012; Soria et al, 2012). However, acquired resistance is unavoidable (Pao et al, 2005). Occurrence of a second EGFR mutation p.T790M in exon 20 represents the most frequent mechanism of acquired resistance(Yu et al, 2013). Osimertinib (AZD9291) is a 3^rd^-generation irreversible EGFR-TKI with potent activities against T790M (Skoulidis & Papadimitrakopoulou, 2017), and has shown significantly higher efficacy in T790M-positive advanced NSCLC patients (Mok et al, 2017). Moreover, osimertinib showed efficacy superior to that of 1^st^ or 2^nd^ generation EGFR-TKIs in the first-line treatment of EGFR mutation-positive advanced NSCLC (Soria et al, 2018). However, it is very disappointing that acquired resistance to such a highly-effective and low-toxicity drug will inevitably occur (Janne et al, 2015). Thus, innovative treatment strategies are urgently needed to fully clarify the mechanisms of acquired resistance to osimertinib.

The mechanisms of osimertinib resistance are diverse and not fully understood. Emerging clinical data suggest that the underlying mechanisms include other EGFR mutations, C797S and L798I, which also prevent drug binding(Chabon et al, 2016; Thress et al, 2015), bypassing of MET or ERBB2 signaling activation(Kim et al, 2015; Mizuuchi et al, 2016; Ortiz-Cuaran et al, 2016; Planchard et al, 2015), or constitutive MAPK pathway activation by mutated KRAS or MEK(Eberlein et al, 2015). Besides, amplification of EGFR wild-type alleles but not mutant alleles is sufficient to confer acquired resistance to osimertinib(Nukaga et al, 2017). However, the majority of patients likely develop resistance by as yet unknown mechanisms. Therefore, it is of great significance to investigate new treatment regimens which can reverse osimertinib resistance caused by diverse mechanism and enhance osimertinib efficacy.

Autophagy is an evolutionarily conserved catabolic process involving the degradation of cytoplasmic constituents, and the recycling of long-lived or aggregated proteins(Yu et al, 2017). In multiple tumor cells, autophagy is upregulated during adverse conditions, including chemoradiotherapy or a nutrient-deficient environment, promoting tumor cell survival; thus, autophagy may be considered a potential mechanism of drug resistance(2014; Auberger & Puissant, 2017; Chen et al, 2016b). In NSCLC treated with EGFR-TKI, autophagy is a double-edged sword contributing to both cell survival and death. Reduced autophagy was related to resistance to erlotinib therapy (Wei et al, 2013). On the other hand, several recent studies have shown that treatment of erlotinib or afatinib induced autophagy and inhibition of autophagy improves the anti-tumor activity of these drugs in lung adenocarcinoma (Hu et al, 2017; Wang et al, 2016). Moreover, the pro-cell survival and pro-cell death roles of autophagy can be switched by adding gefitinib at an early time of hypoxia or by re-activating EGFR at a later time of hypoxia in cancer cell lines (Chen et al, 2016a). Therefore, more works are needed to better understand the role of autophagy in EGFR-targeted therapy for NSCLCs. As a 3^rd^ generation EGFR-TKI, osimertinib has a different chemical structure and different potential resistance mechanisms when compared to 1^st^ or 2^nd^ generation EGFR-TKI. Recently, in osimertinib-sensitive cells, osimertinib was found to induce autophagy (Tang et al, 2017). However, it is unknown whether the accumulated autophagy may induce osimertinib resistance, or whether inhibition of autophagy may restore osimertinib sensitivity.

Therefore, we preformed the current study to clarify the role of autophagy in osimertinib resistance and the potential mechanisms. Since use of osimertinib in 1^st^ or 2^nd^ line may lead to different resistance mechanisms, a series of cell lines were chosen to mimic the clinical usage of osimertinib, including PC-9 cells (19del and sensitive to 1^st^ generation EGFR-TKIs), PC-9GR cells (T790M+ with acquired resistance to the 1^st^ generation EGFR-TKI gefitinib), and H1975 cells (de novo T790M+ with primary resistance to gefitinib), respectively. We first found that osimertinib treatment increased autophagy to a much greater extent than that of gefitinib in osimertinib-sensitive cells, and autophagy inhibitors act synergistically with osimertinib to inhibit cell growth. Next, we demonstrated that enhanced autophagy was a common feature in osimertinib-resistant cells with heterogeneous mutations. Inhibition of autophagy reversed osimertinib resistance. Mechanistically, beclin 1-mediated autophagy determined osimertinib resistance through regulation of stem-cell like properties by upregulating Sox2 and ALDH1, which indeed promote osimertinib resistance. Clinically, enhanced autophagy was also found in several patients with resistance to osimertinib. These findings highlight the importance of beclin 1-mediated autophagy in acquired resistance to osimertinib.

## Results

### Enhanced autophagy lead to drug resistance in osimertinib-sensitive cells

We first interrogated whether osimertinib treatment could induce autophagy and the role of autophagy in osimertinib sensitivity. PC-9GR and PC-9 cells were treated with osimertinib, and then exposed to cyto-ID green detection reagent that selectively labels accumulated autophagic vacuoles. More pre-autophagosomes, autophagosomes, and autolysosomes were observed after osimertinib treatment in PC-9GR and PC-9 cells. Increased LC3 II expression and decreased p62 expression were also found in both cell lines after osimertinib treatment (Fig. 1A and B, Fig. S1). To further confirm whether autophagy was induced after osimertinib treatment, we examined the autophagic flux in osimertinib-treated PC-9 cells using MG132, a potent proteasome inhibitor. Result showed that LC3II was further significantly increased in both cell lines under osimertinib plus MG132 combination treatment compared to osimertinib alone, indicating that osimertinib induced high autophagic flux (Fig. 1B, Fig. S2A). Interestingly, the level of osimertinib-induced autophagy was much higher than that of gefitinib in PC-9 and PC-9GR cells (Fig. 1C, Fig. S2B). Considering previous reports that autophagy play a complex role in gefitinib or erlotinib resistance, we next investigated whether the highly elevated autophagy could affect osimertinib sensitivity. We observed that treatment with SP-1, a specific and potent autophagy inhibitor, abolished osimertinib-induced autophagy increasement and significantly increased osimertinib sensitivity in both cell lines, as determined by the MTT assay (Fig. 1D, Fig. S3). Similar results were obtained with two other autophagy inhibitors, 3-MA (3-Methyladenine, a PI3K inhibitor) and CQ (chloroquine diphosphate salt, which inhibits the integration of autophagosomes with lysosomes) (Fig. S4). The Ki67 incorporation assay revealed that SP-1 combined with osimertinib resulted in a robust inhibition of cell proliferation in PC-9GR cells (Fig. 1E). These results showed that osimertinib treatment induced autophagy in osimertinib-sensitive cells, and inhibition of autophagy increased osimertinib efficacy. We then investigated whether autophagy enhancement by rapamycin, the prototypic inhibitor of mammalian target of rapamycin (mTOR), could decrease osimertinib sensitivity. MTT assay showed that treatment with rapamycin resulted in decreased osimertinib sensitivity (Fig. 1F), and increased autophagy was confirmed by LC3II and p62 level alterations (Fig. 1G, Fig. S5). Taken together, we conclude that autophagy plays an important role in osimertinib sensitivity.

**Figure 1:**
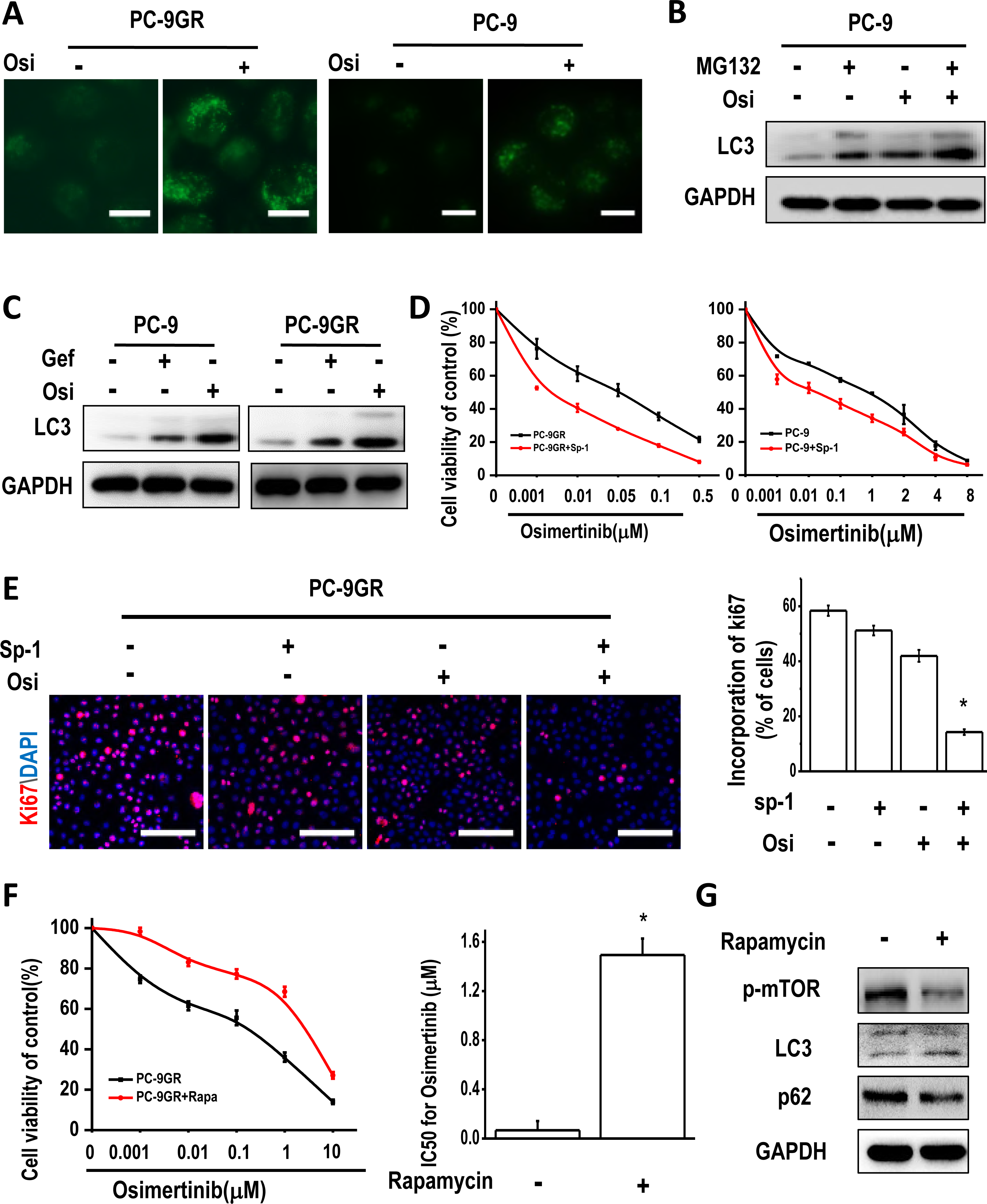
Autophagy inhibition resulted in increased osimertinib sensitivity in osimertinib-sensitive cells. (A) Fluorescent micrographs of autophagosomes in PC-9GR and PC-9 cells treated with or without osimertinib for 24h. Scale bar, 10μm. (B) Western blot showing that high autophagy flux was found in PC-9 cells treated with osimertinib. (C) Osimertinib treatment induced autophagy to a much greater extent than that of gefitinib in both PC-9 cells and PC-9GR cells. Gefitinib, 10nM in PC-9 cells and 4 μM in PC-9GR cells; osimertinib, 20nM in PC-9 cells and 10M in PC-9GR cells. The level of LC3 was examined using Western blot. (D) MTT assay for PC-9GR and PC-9 cells treated with the indicated concentrations of osimertinib for 72h. Experiments were performed in triplicate, and data are mean±SEM. (E) Ki67 staining of PC-9GR cells treated with osimertinib with or without spautin-1(10 μM). Scale bar, 100μm. Experiments were performed in triplicate, and data are mean±SEM. Histogram shows the percentages of Ki67-positive cells in the indicated groups (*p<0.05 by Student’s t test) (F) MTT assay for PC-9GR cells treated with the indicated concentrations of osimertinib with or without rapamycin (500 nM) for 48h. Experiments were performed in triplicate, and data are mean±SEM (*p<0.05 by Student’s t test). (G) Western blot assessment of PC-9GR cells treated with rapamycin for 48h.

### Generation of osimertinib-resistant cell lines

In order to investigated whether autophagy play a role in osimertinib resistance, osimertinib-resistant cell lines were established from PC-9GR cells, H1975 cells and PC-9 cells, respectively, using the single-colony selecting method. The origins of the parental cells were confirmed by the STR loci assay. The osimertinib-resistant cells (PC-9OR, PC-9GROR and H1975-OR cells) displayed similar morphologic features compared to parental cells (Fig. 2A, Fig. S6). Next, the MTT assay was performed to evaluate osimertinib resistance of the cell lines. As shown in Fig. 2B, they were all highly-resistant to osimertinib, with IC_50_ markedly elevated when compared with those of parental cells. Next, the long-term proliferation ability of the resistant cells was evaluated by colony formation assay. Under osimertinib pressure, colonies were found in osimertinib-resistant cells but not in parental cells (Fig. 2C, Fig. S7). These results indicated that these cell lines generated were highly resistant to osimertinib.

**Figure 2:**
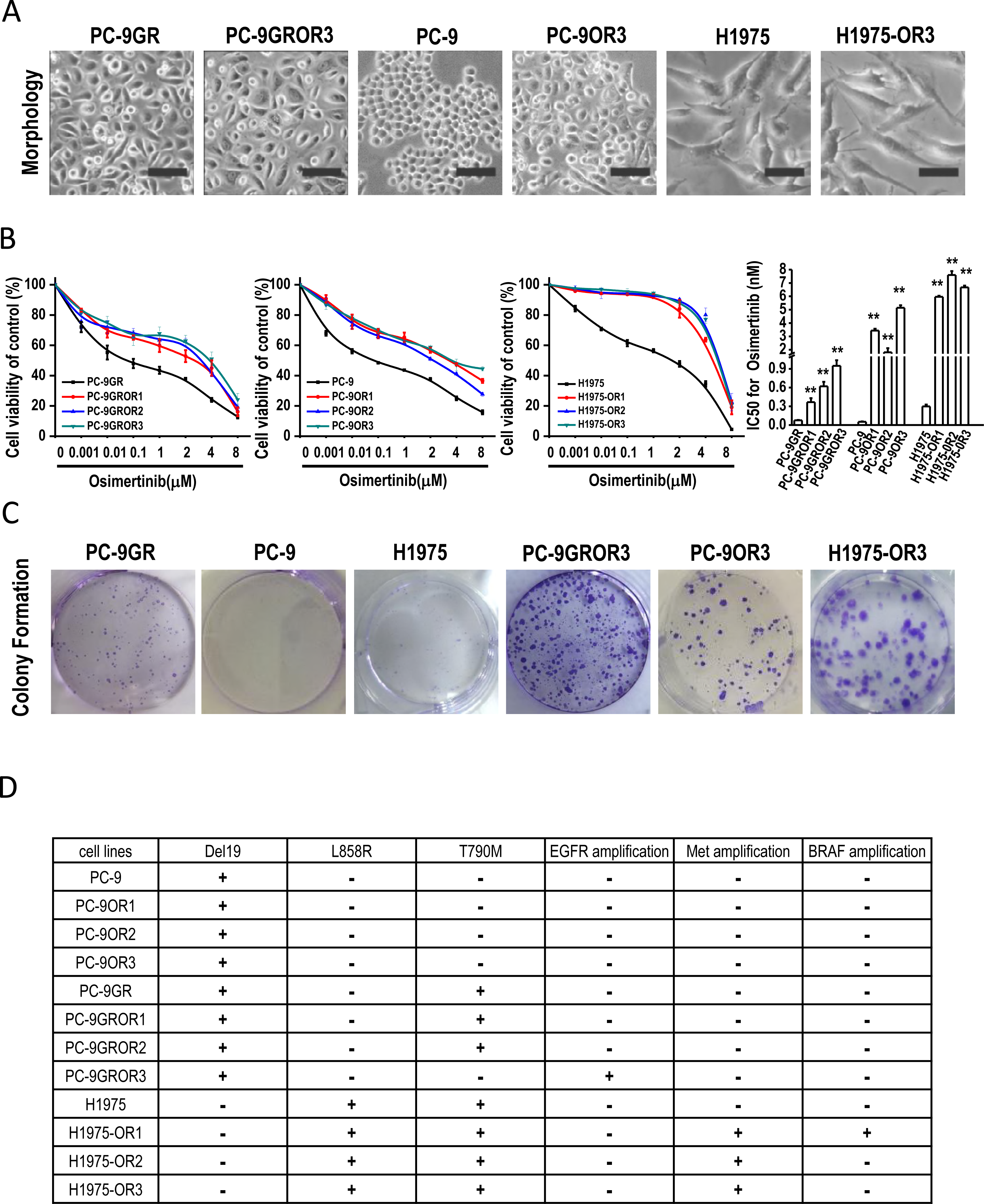
Establishment of osimertinib-resistant cell lines from parental PC-9GR, PC-9, and H1975 cells. (A) Micrographs of parental and the corresponding resistant cells. Scale bar, 30μm. (B) MTT assay for parental PC-9GR, PC-9, H1975 cells and their corresponding resistant cells treated with increasing concentrations of osimertinib for 48 hours. Experiments were performed in triplicate, and data are mean±SEM. Histogram shows IC50 values in the indicated groups (**p<0.01 by Student’s t test). (C) Colony formation assay of resistant and parental PC-9GR, PC-9, and H1975 cell lines. (D) Summary of the gene alterations in each resistant cell lines and parental cell lines detected by whole-exome sequencing.

To clarify potential resistance mechanisms, DNA isolated from parental and resistant clones were subjected to whole exome sequencing. Several genetic alterations potentially relevant to osimertinib resistance were identified (Fig. 2D). *EGFR* amplification and loss of T790M were identified in PC-9GROR3 cells, while *MET* amplification was found in all H1975-OR cells; *BRAF* amplification was detected in H1975-OR1 cells (Fig. 2D). However, no new mutations were found in PC-9OR cells. These results indicated that diverse mechanisms may exist in osimertinib resistance.

### Enhanced autophagy in osimertinib-resistant cell lines determines resistance to osimertinib

We next estimated autophagy levels in osimertinib-resistant cell lines. Increased numbers of accumulated autophagic vacuoles were found in these cells, as shown by fluorescence levels quantified using high-content imaging system (Fig. 3A, Fig. S8). Meanwhile, the protein levels of LC3II was upregulated and p62 was downregulated in PC-9GROR, PC-9OR and H1975-OR cells (Fig. 3B, Fig. S9). Next, we compared autophagy flux in parental and resistant cells. Under MG132 treatment, LC3 II level was highly increased in PC-9OR3 cells when compared with that of parental PC-9 cells (Fig.3C, Fig. S10), indicating high autophagy flux in osimertinib-resistant cells. To further confirm autophagy completion, osimertinib-resistant cells and respective parental cells were assessed by transmission electron microscopy. Autolysosomes, which have a single limiting membrane and contain cytoplasmic/organellar materials at various stages of degradation, can be distinguished from autophagosomes (containing a double limiting membrane) by electron microscopy. More autolysosomes were found in PC-9GROR3, PC-9OR3 and H1975-OR3 cells, compared with respective parental cells (Fig. 2D). We further examined the relative levels of autophagic flux using mCherry-EGFP-LC3 in PC-9GR and PC-9GROR cells. After transfection, autophagosomes were shown as yellow punta (the combination of red and green fluorescence), and autolysosomes were shown as red punta (the extinction of EGFP in the acid environment of lysosomes). As shown in Figure E, both the number of yellow autophagosomes and red autolysosomes (the extinction of EGFP in the acid environment of lysosomes) were increased in the PC-9GROR cells compared to PC-9GR cells. These observations indicated that the autophagic process was completed, rather than blocked at the fusion step.

**Figure 3:**
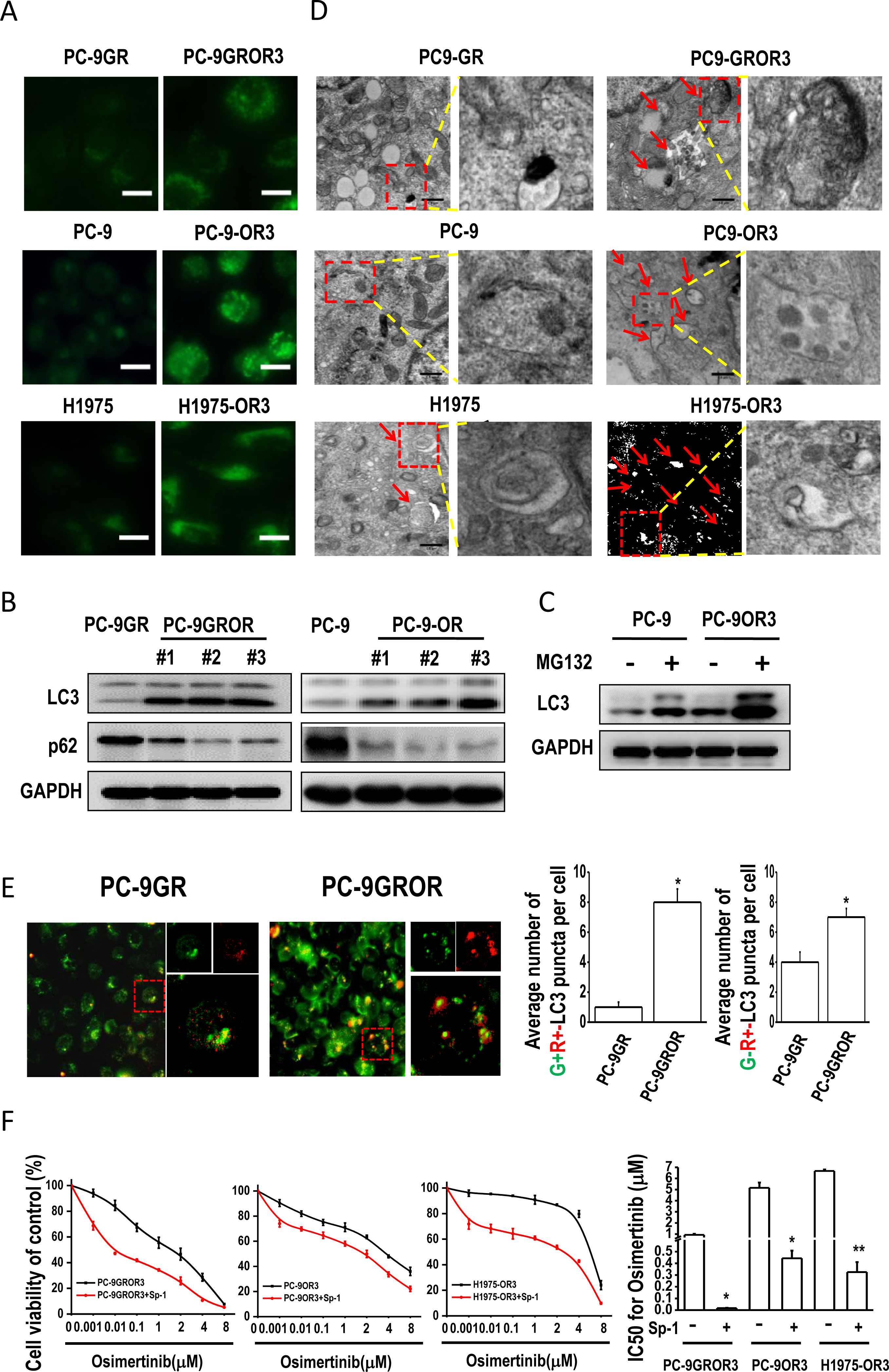
Enhanced autophagy in osimertinib-resistant cells determines osimertinib resistance. (A) Fluorescent micrographs of autophagic vacuoles in parental PC-9GR, PC-9, H1975 cells and the corresponding resistant cells. Scale bar, 10μm. (B) Western blot showing increased LC3-II levels and decreased p62 amounts in resistant cells. (C) MG132 treatment resulted in increased LC3-II levels in resistant PC-9OR3 cells when compared to that of PC-9 cells. (D) Micrographs obtained by transmission electron microscopy showing enhanced autophagosomes in resistant cells compared with parental PC-9GR, PC-9, and H1975 cells. Magnification, 4×10^4^. (E) The level of autophagy flux is increased in PC-9GROR cells. Representative images of mCherry-EGFP-LC3 vector were shown by fluorescent detection. The level of autophagy flux in PC-9GROR cells were increased compared with that in PC-9GR cells. Quantitative analysis of the number of yellow autophagosomesand red autolysosomes. **P < 0.01. (F) MTT assay for parental PC-9GR, PC-9, and H1975 cells and the corresponding resistant cells. Cells were treated with osimertinib with or without autophagy inhibitor, spautin-1 (10μM) for 72 hours. Experiments were performed in triplicate, and data are mean±SEM. Histogram shows IC50 values in the indicated groups (*p<0.05, **p<0.01 by Student’s t test).

Next, we examined the role of enhanced autophagy in osimertinib resistance through pharmacological inhibition of autophagy. SP-1 treatment suppressed autophagy in osimertinib-resistant PC-9GROR, PC-9OR and H1975-OR cells, as shown by decreased LC3 II levels and increased p62 amounts (Fig. S11, 12, 13). Importantly, treatment with SP-1 re-sensitized the resistant cells to osimertinib (Fig. 2E, Fig. S11, 12, 13). Another autophagy inhibitor CQ was also used to treat PC-9OR3 cells, and the cells became more sensitive to osimertinib (Fig. S14). Taken together, autophagy was enhanced in osimertinib-resistant cell lines, and inhibition of autophagy re-sensitized those cells to osimertinib. **Osimertinib-resistant cells have typical stem cell-like properties**

Cancer stem-like cells contribute to tumor heterogeneity and have been implicated in disease relapse and drug resistance (Yeo et al, 2016). To investigated the potential role of stem cell-like properties in osimertinib resistance, pulmosphere formation assay were performed. As expected, PC-9GROR, PC-9OR and H1975-OR cells displayed increased pulmospheres in terms of number and size, comparing to PC-9GR, PC-9 and H1975 cells, respectively (Fig. 4A, Fig. S15). Moreover, CD133 and CD44 are regarded as specific markers for stem-like cells in lung cancer (Nishino et al, 2017; Okudela et al, 2012; Sarvi et al, 2014). FACS analysis revealed that CD133- and CD44-positive cell populations in all osimertinib-resistant cells (PC-9OR, PC-9GROR and H1975-OR) was higher than their parental cells (Fig. 4B). In addition, it was reported that the transcription factor Sox2 and aldehyde dehydrogenase (ALDH) play major roles in stem-like NSCLC cells (Akunuru et al, 2012; Justilien et al, 2014; Sterlacci et al, 2014), either of which is thought to confer drug resistance to tyrosine kinase inhibitors(Dogan et al, 2014; Kim et al, 2014). In our study, higher Sox2 and ALDH1 levels were observed in osimertinib resistant cells when compared to their parental cells (Fig. 4C, Fig. S16). Taken together, these results demonstrated that osimertinib-resistant cells exhibited stem-cell like properties such as enhanced pulmosphere formation ability, higher CD133/CD44 enrichment as well as Sox2 and ALDH1 overexpression.

**Figure 4:**
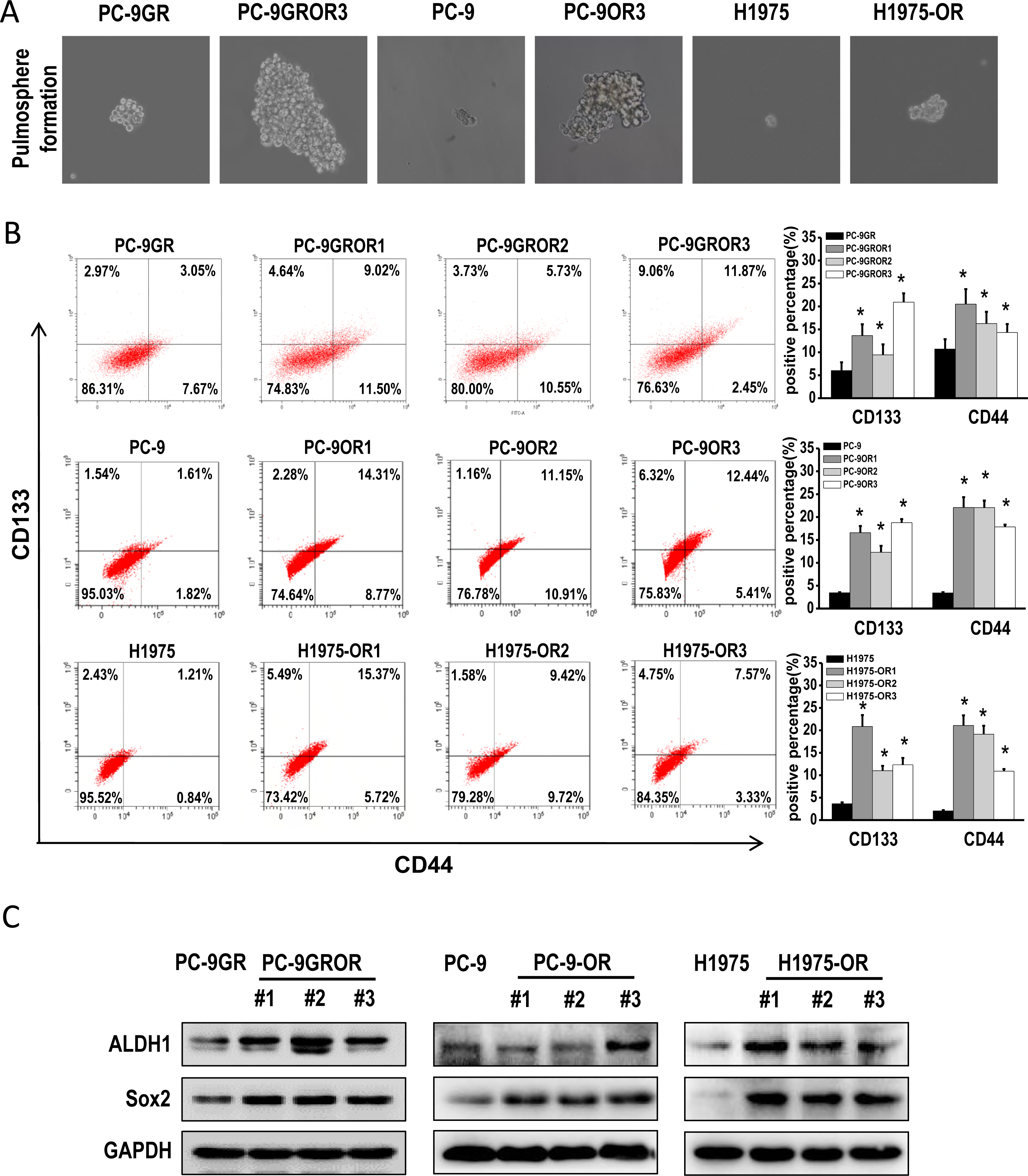
Osimertinib resistant cells show robust stem-cell like properties. (A) Parental and resistant cells were diluted to single cells per well in 6-well low-adhesion plates. Micrographs of spheres formed after 7 days. (B) CD133/CD44 positive cells were detected by flow cytometry using anti-CD133-FITC and CD44-PE antibodies. Top left and top right quadrants represent CD133 positive populations, while top right and lower right are CD44 positive populations. The bar chart shows percentages in various groups (n=3, *p<0.05 by Student’s t test). (C) Western blot showing the expression levels of the potential stem markers ALDH1 and Sox2 in osimertinib-resistant cells.

### Roles of Sox2 and ALDH1 in the maintenance of CSCs and osimertinib resistance

To validate the roles of Sox2 and ALDH1 in stemness, small interfering RNAs targeting *SOX2* and *ALDH1A1* respectively, were designed to investigated the sensitivity of PC-9OR3 cells to osimertinib. First, MTT assay revealed that PC-9OR3 cells were more sensitive to osimertinib after knockdown of either *SOX2* or *ALDH1A1* (Fig. 5A). Secondly, the pulmosphere formation assay showed that *SOX2* or *ALDH1A1* knockdown significantly reduced the number and size of pulmospheres compared with controls (Fig. 5B). Thirdly, colony formation assay demonstrated that *SOX2* or *ALDH1A1* knockdown resulted in significantly decreased clone sizes, suggesting that Sox2 and ALDH1 were responsible for the proliferation of resistant cells (Fig. 5B). Fourthly, PC-9OR3 cells transfected with either *SOX2* or *ALDH1A1* siRNA displayed decreased CD133+ and CD44+ cell populations (Fig. 5C). These findings indicated that Sox2 and ALDH1 might be essential for maintaining stemness and resistance to osimertinib. In addition, ALDH1 protein levels decreased after silencing of *SOX2*, whereas no Sox2 protein levels change was observed after *ALDH1A1* knockdown (Fig. 5D, Fig. S17). These findings demonstrated that the role of ALDH1 in maintaining stemness and osimertinib resistance might be mediated by Sox2.

**Figure 5:**
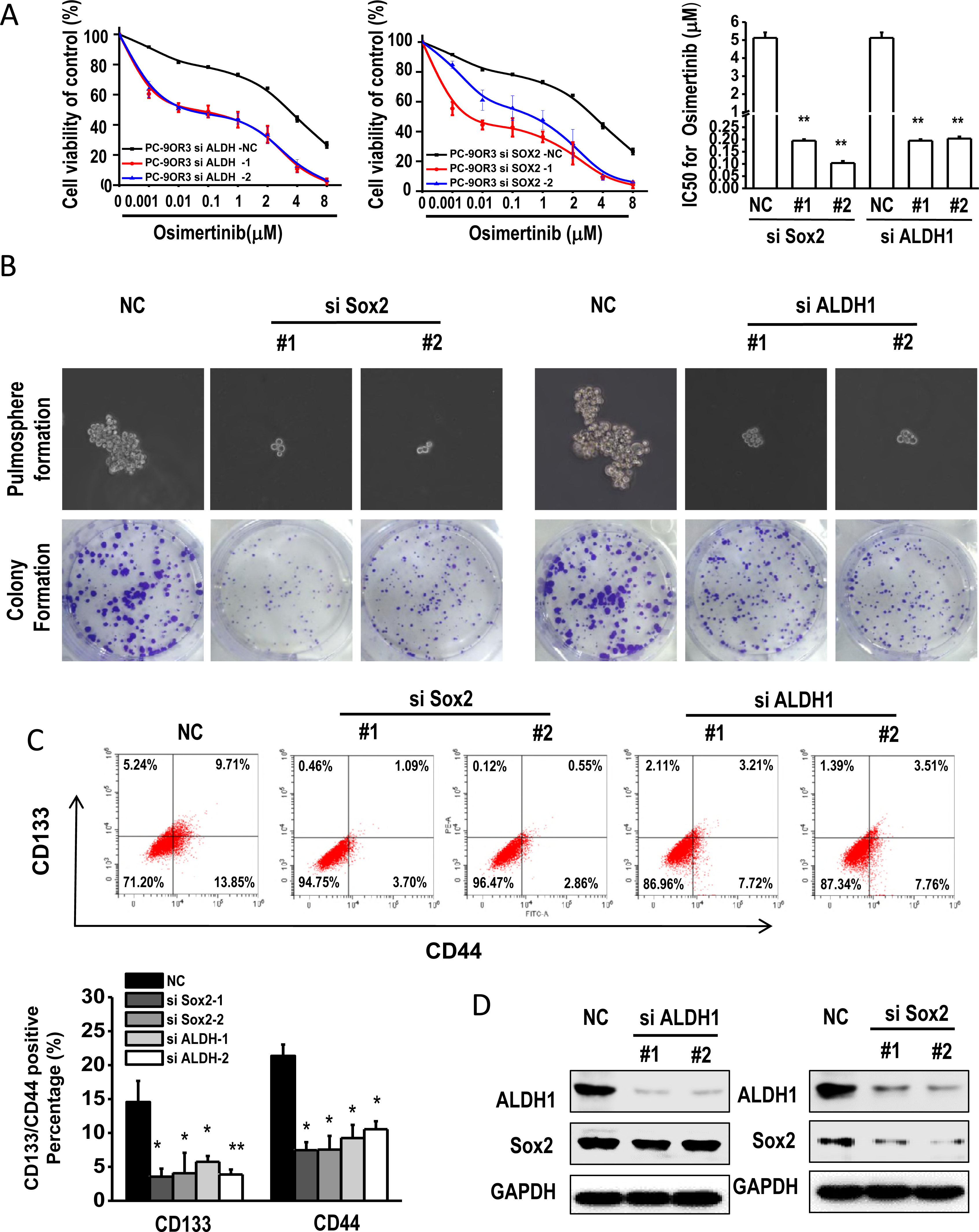
Loss of Sox2 and ALDH1 affects stemness and osimertinib resistance. (A) MTT assay for PC-9OR3 cells transfected with control, Sox2, and ALDH1A1 siRNAs, respectively, treated with increasing concentrations of osimertinib for 48 hours. Experiments were performed in triplicate, and data are mean±SEM. Histogram shows IC50 values in the indicated groups (**p<0.01 by Student’s t test). (B) Colony formation and pulmosphere formation assays for assessing the PC-9OR3 cell line after transfection with control, Sox2, and ALDH1A1 siRNAs, respectively. (C) CD133/CD44 positive cells were detected by flow cytometry with anti-CD133-FITC and CD44-PE antibodies. The bar chart shows percentages in various groups (n=3, p<0.05 by Student’s t test). (D) Western blot showing the expression levels of Sox2 and ALDH1 in the PC-9OR3 cell line after transfection with ALDH1A1 and Sox2 siRNAs, respectively.

### Beclin 1-dependent, not Atg5-related autophagy maintains stem-like cell properties in osimertinib-resistant cells

We next investigated the mechanism of autophagy induced osimertinib resistance by maintaining stem cell-like properties. SP-1 treatment lead to smaller spheres PC-9OR3 cells, compared to control (Fig. 6A and Fig. S18). Flow cytometry also revealed that SP-1 resulted in decreased population rates of CD133 and CD44 positive cells (Fig. 6B, Fig. S19). These findings indicated autophagy inhibition can decrease stem cell-like characteristics in osimertinib-resistant cells. In addition, we found that SP-1 treatment downregulated ALDH1 and Sox2 in osimertinib-resistant cells (PC-9OR1, 2, 3) (Fig. 6C, Fig. S20). Similar observations were obtained in PC-9GROR3 cells and H1975-OR3 cells (Fig. S21). These results demonstrated that autophagy inhibition can result in decreased stemness in osimertinib-resistant cells.

**Figure 6:**
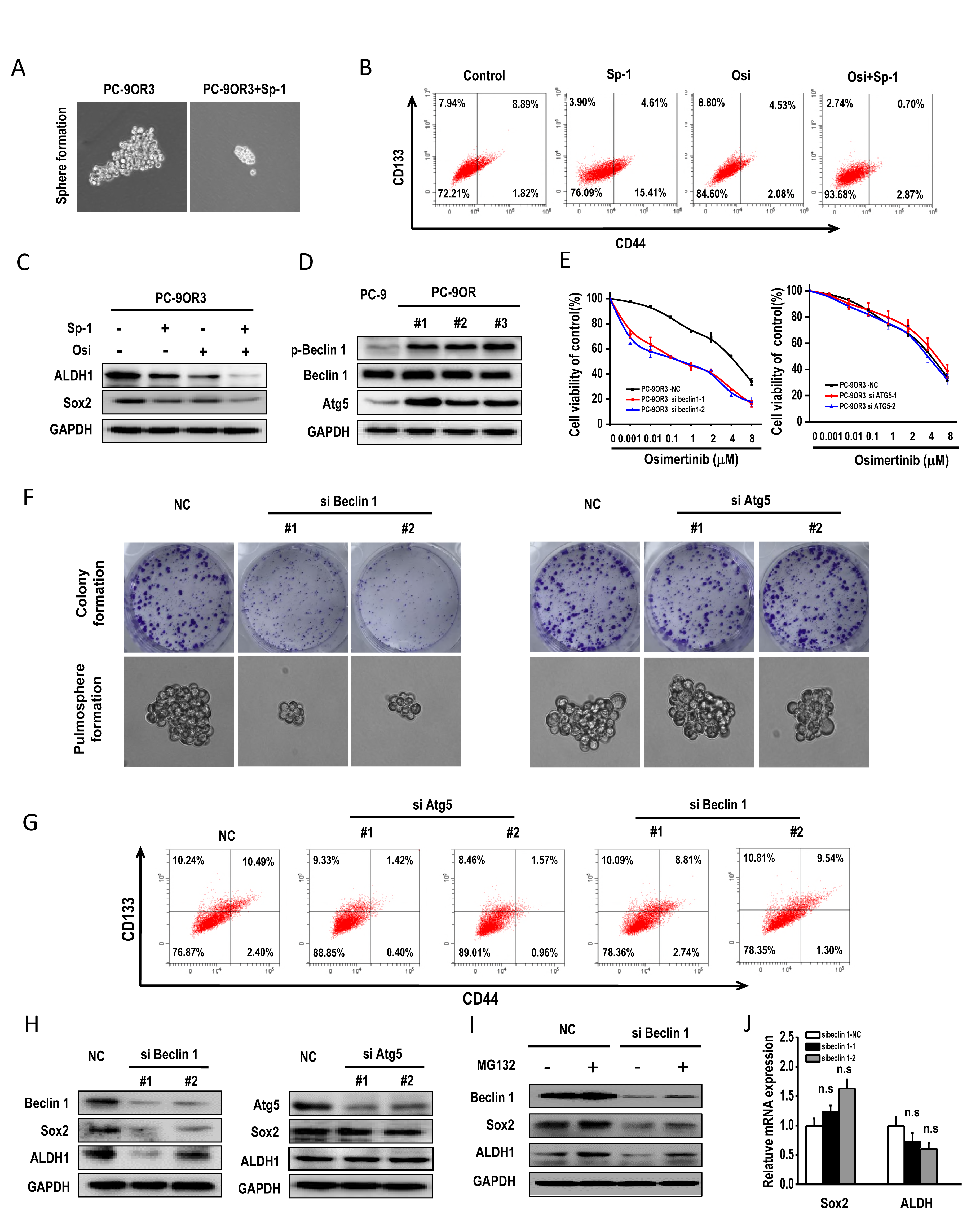
Beclin 1-mediated, not Atg5-dependent autophagy is critical for stem-cell like properties and osimertinib resistance. (A) Pulmosphere formation assay for PC-9OR3 cells treated with DMSO or the autophagy inhibitor spautin-1 (10μM). (B) CD133/CD44 positive PC-9OR3 cells were detected after treatment with spautin-1 alone or combined with osimertinib. (C) ALDH1A1 and Sox2 levels measured by Western blot in PC-9OR3 cells after treatment with spautin-1 and osimertinib. (D) Phospho-beclin 1 (Ser93), total beclin1, and Atg5 levels measured by Western blot in parental PC-9 and resistant PC-9OR3 cells. (E) Cell viability of PC-9OR3 cells transfected with control, beclin 1, and Atg5 siRNA, respectively. Experiments were performed in triplicate, and data are mean±SEM. (F) Colony formation and pulmosphere formation assays for PC-9OR3 cells after transfection with control, beclin 1, and Atg5 siRNAs, respectively. (G) CD133/CD44 positive PC-9OR3 cells were detected after transfection with control, beclin 1, and Atg5 siRNAs, respectively, by flow cytometry with anti-CD133-PITC and CD44-PE antibodies. (H) Sox2 and ALDH1 levels were measured in PC-9OR3 cells after transfection with control, beclin 1, and Atg5 siRNAs, respectively. (I) Sox2 and ALDH1 levels were measured in PC-9OR3 cells after transfection with control or beclin 1siRNAs following with MG132 treatment for 6h. (J) qPCR analysis of mRNA level of Sox2 and ALDH1 after beclin 1 knockdown.

Atg5 is an essential gene in canonical macroautophagy, while the non-canonical autophagic pathway, which is independent of Atg5, has been reportede (Honda et al, 2014; Ma et al, 2015). Next, we investigated whether Atg5-dependent autophagy maintains stem cell-like characteristics. Atg5 and phosphorylated beclin 1 (Ser 93) levels were increased in resistant cells, while total Beclin 1 expression remained unchanged (Fig. 6D and Fig.S22). Similar results were found in other osimertinib resistant cells (Fig. S23). Treatment with SP-1 resulted in decreased beclin 1 phosphorylation but not total beclin 1 and Atg5 amounts (Fig. S24). We silenced Atg5 and beclin 1 by siRNAs to examine their effects on osimertinib resistance. Knockdown of beclin 1 resulted in enhanced sensitivity of PC-9OR3 cells to osimertinib, whereas Atg5 knockdown showed no remarkable effects (Fig. 6E, Fig. S25). Furthermore, colony and pulmosphere formation assays demonstrated that beclin 1, not Atg5, was essential for stem cell-like properties, as decreased colony formation and smaller pulmospheres were observed only in beclin 1 knockdown cells (Fig. 6F). In addition, siRNA targeting beclin 1 led to a significant decrease of CD133/CD44-positive cells while siRNA targeting Atg5 showed no significant effects (Fig. 6G, Fig. S26).

Next, the effects of beclin 1 knockdown on Sox2 and ALDH1 protein levels were evaluated. Results showed that beclin 1 knockdown resulted in decreased Sox2 and ALDH1 protein amounts (Fig. 6H, Fig. S27), and beclin 1 knockdown also weakened the accumulation of Sox2 and ALDH1 proteins after the proteasome inhibitor MG132 treatment (Fig. 6I, Fig. S28). Interestingly, mRNA levels of *SOX2* and *ALDH1A1* were unchanged after beclin 1 knockdown (Fig. 6J). This suggested that beclin1, but not Atg5, might maintain stemness through preventing the protein degradation of Sox2 and ALDH in osimertinib-resistant cells.

### Autophagy inhibition enhances the anti-tumor activity of osimertinib in PC-9GR/mouse xenografts

We next assessed whether combination of the autophagy inhibitor CQ and osimertinib is more effective in xenografts established with PC-9GR cells. Result showed that CQ treatment slightly reduced tumor growth in PC-9GR xenografts, and osimertinib alone resulted in significant tumor shrinkage. The combination of CQ and osimertinib can further inhibit tumor growth (P < 0.05 compared with osimertinib alone; Fig. 7A). During the treatment, no overt weight loss was observed in mice treated with CQ and/or osimertinib (Fig. 7B). Collectively, these findings suggested that CQ enhanced the therapeutic efficacy of osimertinib *in vivo*.

**Figure 7:**
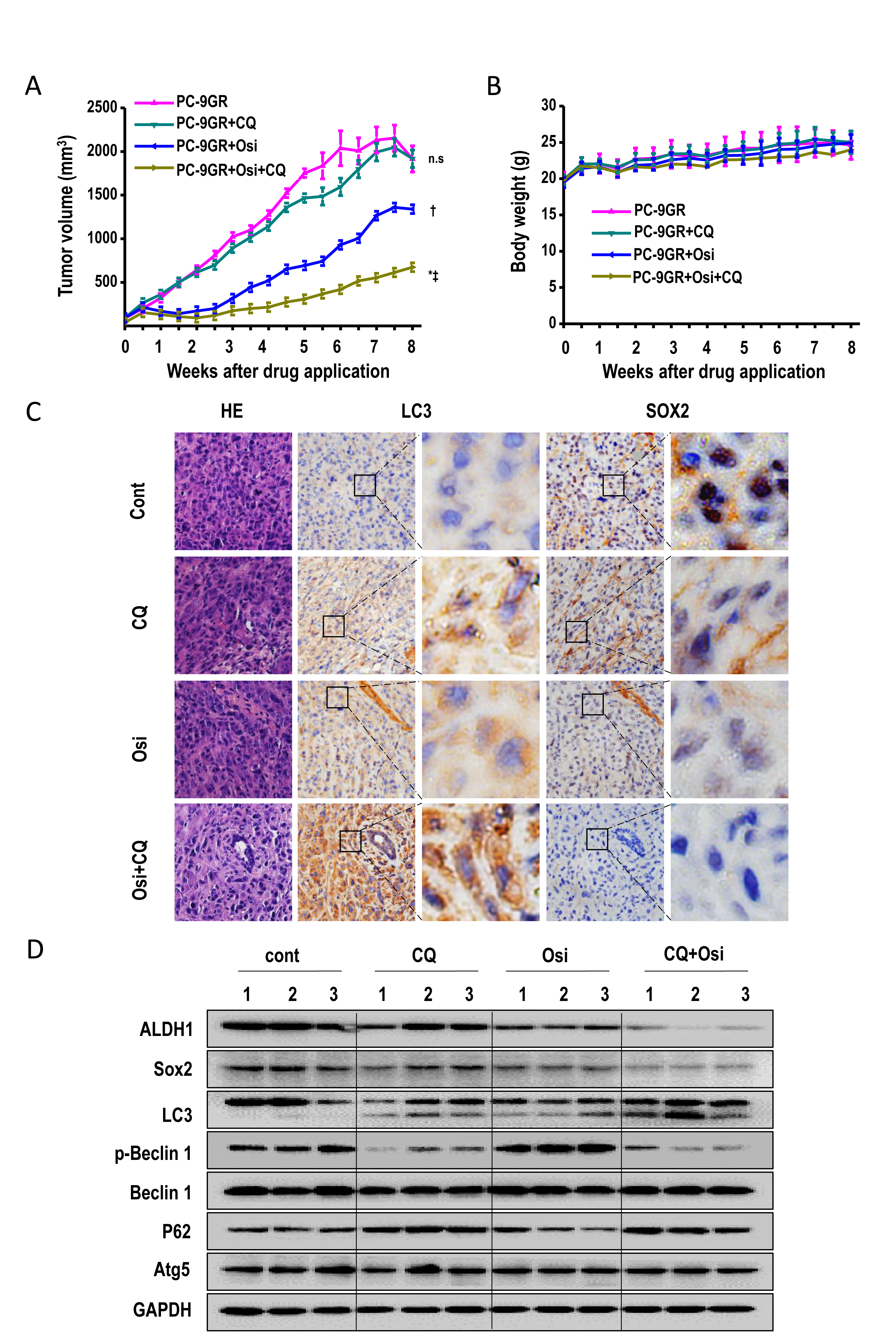
Combination of the autophagy inhibitor CQ and osimertinib effectively inhibits the growth of PC-9GR xenografts. (A) PC-9GR xenografts were treated with control, CQ, osimertinib, and combined CQ/osimertinib, for 8 weeks. Tumor sizes were presented as mean±SEM (n=8); n.s, not significant compared with the control group; *, P<0.001 compared with the control group; †, P<0.01 compared with the control group; ‡, P<0.05 compared with the osimertinib alone group. (B) Body weight were presented as mean±SEM (n=8); n.s, not significant. (C) Representative immunohistochemical staining results for LC3 and Sox2, and hematoxylin-eosin staining for tumor xenografts from nude mice. (D) Whole protein cell lysates were prepared randomly from 3 tumors per group for Western blot to detect the indicated proteins.

Next we explored the mechanism of combined therapy which was more effective than monotherapy in PC-9GR xenografts. Immunohistochemical staining showed high expression of LC3 and low Sox2 levels in the combination group (Fig. 7C). Osimertinib treatment resulted in increased Beclin-1 phosphorylation, while the combination therapy decreased Beclin-1 and Sox2 phosphorylation (Fig. 7D). These findings suggested that CQ/osimertinib combination was associated with the inhibition of autophagy and stem cell like properties *in vivo*.

### Enhanced autophagy was found in NSCLC patients with resistance to osimertinib

Next, we investigated whether enhanced autophagy existed in NSCLC patients with resistance to osimeritnib. A retrospective analysis was performed by enrolling 39 NSCLC patients who had developed drug resistance to osimertinib from August 2015 to Feb 2018 in our hospital. Prior to treatment of osimertinib, all these patients displayed resistance to 1st-generation EGFR-TKI and *EGFR* T790M mutation was detected in them. After osimertinib resistance, either plasma or tissue biopsies from these 39 patients were profiled by capture-based targeted ultra-deep sequencing. As shown in Fig. 8A, a series of potential resistance mechanisms were found, including EGFR C797S mutation, MET amplification, ERBB2 amplification, KRAS mutation, PI3K mutation, et al. Of note, 57% patients developed resistance with unknown mechanisms. We next examined LC3 expression in 5 patients with paired tumor tissue samples (before osimertinib treatment and after osimertinib resistance). Before osimertinib treatment, low LC3 expression was found in all 5 patients (Fig. 8B). After osimertinib resistance, elevated LC3 expression was found in 3 patients (Patient #1, 4 and 5). Overall mutation spectrum of the 5 patients was displayed in Fig. 8C. Of the 3 patients with increased LC3 expression after osimertinib resistance, EGFR C797S mutation was found in 1 patient, sensitive EGFR mutations were found in the other 2 patients. In the remaining 2 patients without LC3 level increasing, C-met amplification was identified. Taken together, these results indicate that enhanced autophagy exist in at least some NSCLC patients with resistance to osimertinib.

**Figure 8:**
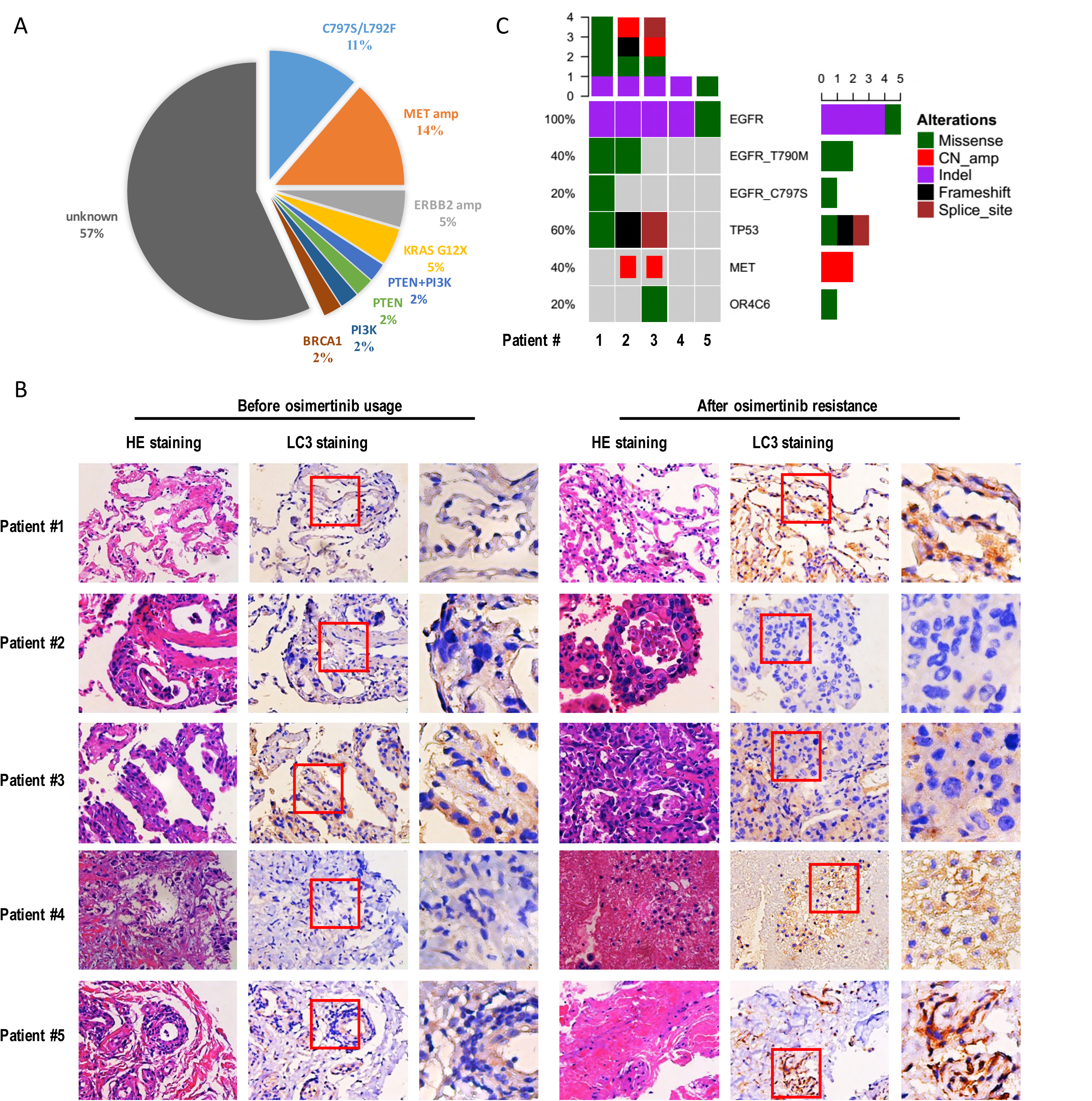
Enhanced autophagy was found in patients with acquired resistance to osimertinib. (A) A chart showing potential resistance mechanisms to osimertinib in 39 NSCLC patients. More than 50% patietns developed resistance by yet unknown mechanisms. (B) Immunohistochemical staining results for LC3 and hematoxylin-eosin staining in paired tumor sections from 5 patients (before osimertinib treatment and after osimertinib resistance). Positive staining was seen in patient #1,4 and 5 after osiemrtinib treatment. (C) Overall mutation spectrum of the 5 patients. Different color presents different types of baseline mutation. The top bar demonstrated the number of mutations detected in an individual patient. The side bar stands for the number of patients harboring the corresponding mutation.

## Discussion

Currently there is no effective approach to overcome acquired resistance to 3^rd^-generation EGFR-TKI osimertinib. The current study demonstrated that enhanced autophagy not only induced drug resistance in osimertinib-sensitive cells, but also was a general feature in osimertinib-resistant cells which presents diverse and heterogeneous mutations. Autophagy inhibitors and osimertinib synergistically inhibited the growth of both sensitive and resistant tumor cells. Enhanced stem-cell like properties were found in osimertinib-resistant cells. Of note, beclin 1-mediated autophagy helped maintain stem cell-like properties by upregulating Sox2 and ALDH1, which indeed facilitate osimertinib resistance. CQ in combination with osimertinib significantly inhibited tumor growth in xenograft experiments. Taken together, we have shown that pro-survival autophagy determines osimertinib resistance through regulation of stem-cell like properties.

Role of autophagy in lung cancer targeted therapy is perplexing. In advanced lung adenocarcinoma treated with gefitinib, ATG5 rs510532 and ATG10 rs10036653 genetic variations in autophagy core genes are significantly associated with clinical outcomes (Yuan et al, 2017). Previously, reduced autophagy was related to resistance to erlotinib therapy (Wei et al, 2013), and when autophagy is further elevated by a treatment in addition to 2^nd^ generation EGFR-TKI afatinib, it can induce autophagic cell death (Lee et al, 2015). On the other hand, reports demonstrated that gefitinib and erlotinib induced pro-cell survival autophagy in both sensitive and resistant cancer cells (Han et al, 2011; Sugita et al, 2015; Zou et al, 2013). Combining glucose deprivation and autophagy inhibitor could synergize and overcome the acquired resistance against erlotinib (Ye et al, 2017). Taken together, the role of autophagy in resistance to 1^st^ and 2^nd^ generation EGFR-TKI is contradictory. The chemical structure of osimertinib is totally different from 1^st^ and 2^nd^ generation EGFR-TKI, and the role of autophagy in osimertinib resistance is unknown.

In the current study, we found a striking difference between gefitinib and osimertinib-induced autophagy. In gefitinib-resistant PC-9GR cells, the level of autophagy was only slightly higher than that of parental PC-9 cells, while a much higher level of autophagy was found in osimertinib-resistant cell lines than their parental cells. Moreover, in both PC-9 cells and PC-9GR cells, osimertinib induced autophagy to a much greater extent than that of gefitinib. Significantly, inhibition of autophagy by several inhibitors and si-RNAs *in vitro* resulted in enhanced osimertinib efficacy, and the combination of CQ and osimertinib *in vivo* markedly decreased tumor growth than osimertinib alone. Clinically, enhanced autophagy was found in several patients with resistance to osimertinib. These results indicate that pro-cell survival autophagy leads to osimertinib resistance. Previously, activation of pro-survival autophagy has been found in therapeutics of many cancer, and blockage of autophagy promotes cell death (Amrein et al, 2011; Han et al, 2008; Lee et al, 2017). Inhibition of autophagy has been proposed as a new approach to enhance efficacy of targeted therapy. For example, simultaneously targeting Hedgehog signaling pathway and autophagy could overcome drug resistance of BCRABL-positive chronic myeloid leukemia to imatinib (Zeng et al, 2015). Elevated autophagy activity contributes to the enhanced tolerance to metabolic stresses of EGFRvIII-expressing cells in glioblastoma. Targeting this survival mechanism abrogates this advantage and results in enhanced tumor cell killing (Jutten et al, 2018). Taken together, our results with those findings suggest that pro-cell survival autophagy plays an important role in targeted therapy of cancer.

Targeting autophagy may be developed as a new approach to overcome osimertinib resistance clinically. The current study established osimertinib-resistant cell lines from PC-9 cells, which have only a sensitive EGFR mutation, and PC-9GR and H1975 cells, in which T790M is present. This choice of cells reflected the clinical application of osimertinib in 1^st^ or 2^nd^ line settings. Besides, loss of T790M, EGFR amplification, Met amplification, and BRAF amplification were found in osimertinib-resistant cell lines, in line with the clinical situation that diverse mutations of known driver genes and unknown mechanisms faced by patients with osimertinib resistance (as shown in Fig. 8A). Interestingly, enhanced autophagy were found in all resistant cell lines and several patients with different potential resistance mechanisms to osimertinib, which indicates that autophagy inhibition may be effective in osimertinib-resistant patients with heterogeneous resistance mechanisms. In the current study, we found that CQ in combination with osimertinib *in vivo* markedly decreased tumor growth. CQ is an FDA approved drug used for malaria, rheumatoid arthritis, and other autoimmune diseases, and is very cheap with an established history of good tolerability. Therefore, CQ may be applied together with osimertinib clinically to enhance osimertinib efficacy or to overcome osimertinib resistance.

The underlying mechanism of how autophagy renders osimertinib resistance is unknown. The importance of stemness in tumor heterogeneity and the heterogeneity of resistance mechanisms found in osimertinib-resistant cell lines in the current study initiated us to investigate whether autophagy may regulate osimertinib resistance through regulation of stem cell-like properties. In fact, osimertinib-resistant cells exhibited stem-cell like properties of enhanced pulmosphere formation ability, high CD133/CD44 enrichment as well as Sox2 and ALDH1 overexpression. Moreover, siRNA knockdown of *SOX2* or *ALDH1A1* increased osimertinib sensitivity, decreased CD133+ and CD44+ populations as well as pulmosphere formation ability. These results indicate that Sox2-mediated ALDH1 expression was involved in maintaining stemness and conferring osimertinib resistance. Previously, stem cell-like features, including overexpression of putative stem cell markers ALDH1A1 and ABCB1, were observed in cells with acquired resistance to gefitinib or afatinib (Hashida et al, 2015; Shien et al, 2013). Also, stem cell-like characteristics were found in gefitinib-resistant cells, and knockdown of IL-8 led to loss of stem cell-like characteristics and enhanced gefitinib sensitivity (Liu et al, 2015). Therefore, our results, together with previous findings, indicate that enhanced stem cell-like properties mediate osimertinib resistance.

We next asked whether autophagy controls osimertinib resistance through regulation of stem cell-like properties. Previously, it was reported that autophagy suppresses hematopoietic stem cell metabolism by clearing active, healthy mitochondria to maintain stemness (Ho et al, 2017). Here, we addressed the importance of Beclin 1 in maintaining stem-cell like properties. We found that beclin 1 knockdown by siRNA resulted in complete suppression of stem-cell like properties (decreased formation of pulmospheres and reduced levels of the stem cell markers Sox2, ALDH1, and CD133/CD44), which are associated with osimertinib resistance. We also demonstrated that beclin 1 help prevent the protein degradation of Sox2 and ALDH to maintain stemness. These findings support a new physiological role for Beclin 1-dependent alternative macroautophagy in stem-like cell maintenance. Therefore, we hypothesized that Beclin 1 is beneficial for the maintenance of cancer stem-like cells by preventing the protein degradation of Sox2 and ALDH1.

Several studies have reported the role of autophagy in control of stemness of cancer cells. Autophagy maintains the stemness of ovarian cancer stem cells through regulation of FOXA2 (Peng et al, 2017), and inhibition of autophagy reduces chemoresistance and tumorigenic potential of ovarian cancer stem cells (Pagotto et al, 2017). Autophagy promotes the formation of vasculogenic mimicry by glioma stem cells through induction of KDR/VEGFR-2 activation (Wu et al, 2017). In acute myeloid leukemia stem cells, autophagy confers resistance to BET inhibitor JQ1 (Jang et al, 2017). Overall, these reports together with findings of the current study indicate that autophagy has a key role in maintenance of stemness of cancer cells, which then contribute to therapeutic resistance.

Since autophagy is important for osimertinib resistance as shown above, it is reasonable to ask which autophagy genes are involved. Canonical autophagy is mediated by evolutionarily conserved autophagy-related genes (Atg genes), among which Atg5 is considered an essential component(Kim et al, 2013). Recently, Atg5-independent autophagy was reported. Both canonical and Atg5-independent non-canonical autophagic pathways have the same upstream autophagy initiation mechanism, regulated by several autophagic proteins, including Unc-51-like kinase 1 (Ulk1) and Beclin 1. Although higher expression levels of p-Beclin1 (Ser93) and Atg5 were observed in all osimertinib-resistant cell lines, osimertinib resistance was indeed inhibited by beclin 1 knockdown but not Atg5 silencing. This study firstly showed that Beclin 1-dependent and Atg5-independent alternative macroautophagy mediated osimertinib resistance.

## Conclusion

In summary, this study delineates a previously unknown function of autophagy in promoting stemness and osimertinib resistance. Such findings are critical for devising a potential therapeutic strategy to overcome osimertinib resistance. In the future, more clinical work are needed to study whether autophagy level was enhanced in osimertinib-resistant patients and to test the efficacy of autophagy inhibition in combination with osimertinib in EGFR-mutant patients.

## Materials and Methods

### Cell lines

PC-9 cells and gefitinib-resistant PC-9GR cells were generously provided by Prof. J. Xu and Dr. M. Liu (Guangzhou Medical University, China). H1975 cells were from the American Type Culture Collection (ATCC). To establish osimertinib-resistant cell lines, the parental cells were treated with osimertinib at the concentration of IC_50_ for 2 weeks, with higher drug levels for another 3 weeks. The latter dosage was sufficient to kill all parental cells. When resistant clones were visible, the cells were diluted to a single cell per well, and continuous culture was performed in presence of osimertinib. All cells were cultured in RPMI-1640 (Hyclone) with Earle’s salts, supplemented with 10% FBS (Gibco), 2 mmol/L L-glutamine (Gibco), 100U/ml penicillin (HyClone), and 100µg/mL streptomycin (Hyclone) at 37°C, with 5% CO_2_ and 90% humidity.

### Reagents

Osimertinib (TAGRISSO) was obtained from Astra Zeneca. Spautin-1 (S7888), 3-Methyladenine (S2767) and rapamycin (S1039) were purchased from Selleck. Cycloheximide (C6628) was purchased from Sigma-Aldrich, and cycloheximide (HY-12320) from MedChem Express. Anti-LC3II (#12741S), SQSTM1/p62 (#8025S), Atg5 (#9980S), Beclin 1 (#3495S), phospho-(Ser93)-beclin1 (#14717S), Sox2 (#3579S), ALDH1A1 (#36671S), GAPDH (#2118S) antibodies were from Cell Signaling Technology.

### Cell growth assays

The MTT cell proliferation assay was performed as previously described(Yao et al, 2010). Briefly, 2×10^3^ cells/well were seeded in 96-well plates and treated with osimertinib or dimethyl sulfoxide (DMSO) 24 hours later. Absorbance was measured 72 hours after treatment. All experiments were repeated for at least three times. Cell proliferation was also assessed by the Ki67 incorporation assay with a Ki67 labeling and detection kit (BM2889, Boster). Briefly, cells were treated with osimertinib for 48h, incubated for 6h with Ki67 (1:200 dilution), and fixed. Cells were counterstained with 4′, 6-diamidino-2-phenylindole (DAPI) and observed under a fluorescence microscope.

### Colony-formation assay

Briefly, 500 cells were resuspended in culture medium and seeded in six-well plates. After 14 days of culture, the cells were fixed with 4% paraformaldehyde and stained with 0.1% crystal violet. Colonies with a diameter greater than 1 mm were counted. Triplicate samples were used in the experiment.

### Transmission electron microscopy (TEM)

The cells were pre-fixed with 2.5% glutaraldehyde in 0.1M PBS (pH 7.4) for 2h at room temperature, and post-fixed with 1% osmium tetroxide for 2h. The samples were then dehydrated in increasing concentrations of ethanol (50%, 70% and 100%) and acetone, and finally embedded in Araldite. Fifty to sixty nanometer sections were cut on a LKB-I ultramicrotome and transferred to copper grids, post-stained with uranyl acetate and lead citrate, and examined by Gatan JEM-1400 plus transmission electron microscopy.

### Whole-exome sequencing

Whole-exome sequencing libraries were prepared with 3 mg DNA. Exomes were captured using the NimbleGen SeqCap Non-Standard Material 110823-HG19-BEx-L2R-D03-EZ for whole exome sequencing, and libraries were hybridized to custom-designed biotinylated oligonucleotide probes (Roche NimbleGen, USA) covering the target region sequence for target-capture sequencing. DNA sequencing was carried out on a HiSeq Sequencing System (Illumina, CA) with 2×151-bp and 2×76-bp paired-end reads for WES and target-capture sequencing, respectively. Raw sequencing reads were filtered to obtain clean reads, which were then aligned to human genome assembly HG19 with Burrows-Wheeler Aligner (BWA) (Newman et al, 2016). Reads with multiple mapping loci in the genome, and those with more than three mismatches, more than one gap, or a gap of more than 20 base long were removed. Reads harboring an Indel within 5 bp of the fragment ends were removed. Duplicated reads derived from PCR amplification were marked with Picard tools (http://broadinstitute.github.io/picard/). Local realignments and base-quality recalibrations were performed with the GATK software (https://www.broadinstitute.org/gatk/).

### Measurement of autophagic activity and autophagy flux

Autophagic activity was monitored with the Cell-ID Green Autophagy Detection Kit (Enzo Life Sciences, France). The Cell-ID Green autophagy dye serves as a selective marker of autolysosomes and early autophagic compartments. Cells were trypsinized, washed with Assay Buffer, and incubated with Cell-ID Green Detection Reagent for 30 min at room temperature, according to the manufacturer’s instructions. Afterwards, 10,000 events/sample were analyzed by fluorescence microscopy. The 24-well plates were imaged on an ImageXpress Micro (Molecular Devices) high-throughput imager. Image analysis was performed with the MetaXpress software.

To measure autophagy flux, pBABE-EGFP-mCherry-MAP1LC3B (22418, deposited by Jayanta Debnath) plasmid was obtained from Addgene and detailed methods was described previously (Gump et al, 2014). PC-9GR and PC-9GROR cells were transduced with this plasmid for 6h and then replaced fresh media. We used ImageXpress Micro XLS Widefield High-Content screening System to automatic scanning the fluorescent dots of autophagosomes labeled mCherry-EGFP-LC3 after an interval 12h., and the final time point was 72h. Autophagic flux was determinedby the yellow punta (the combination of red and green fluorescence), and red punta (the extinction of EGFP in the acid environment of lysosomes).

### Western blot

Cells harvested by scraping were washed twice with PBS and lysed for 30 min at 4°C in RIPA buffer (Sigma-Aldrich, France). After centrifugation at 12000×g for 15 min at 4°C, the protein content was determined by the BCA assay. Equal amounts of protein were submitted to gel electrophoresis for 2h at 110 V, followed by transfer onto PVDF membranes (90 min, 200 mA) (Roche, Switzerland). Membranes were blocked with 5% bovine serum albumin (BSA) for 1h at room temperature and incubated overnight at 4°C with primary antibodies. Subsequently, the membranes were washed and incubated with 0.02 µg/ml horseradish peroxidase (HRP)-conjugated goat anti-rabbit or anti-mouse IgG (Cell Signaling Technology, USA) for 1h, and visualized with ChemiDoc Touch System (Bio-Rad, USA).

### Pulmosphere formation assay

For tumor pulmosphere formation, 1×10^3^ tumor cells were seeded in non-adhesive 6-well plates. One week after seeding, cell spheres characterized by tight, spherical, non-adherent colonies > 90μm in diameter were observed. All experiments were repeated at least three times.

### Flow cytometry

For CD133 and CD44 staining, 10^6^ cells were incubated with 10μl of each anti-CD133-PE (AC133 clone; Miltenyi Biotech) and anti-CD44-FITC (REA690 clone; Miltenyi Biotech) antibodies diluted in 100μl of staining solution for 15 min at 4°C. Then, 400μl buffer was added, and samples were analyzed with the CytExpert software (Beckman Coulter, USA).

### siRNA Transfection

Small interfering RNAs (siRNAs) were synthesized by Shanghai GenePharma Co., Ltd. (Shanghai, China). For efficacy evaluation, 80 pmol siRNA and negative control siRNA (siNC), respectively, were transfected into PC-9OR cells cultured in 6-well plates using Lipofectamine RNAiMAX, following the manufacturer’s instructions. At 72h post-transfection, knockdown efficiency was determined by examining endogenous expression by Western blot.

### Quantitative RT-PCR

Total RNA was isolated from cultured cells with TRIzol reagent (#15596026, Thermo Scientific, USA), and subjected to reverse transcription with PrimeScript RT reagent Kit containing gDNA Eraser (TAKARA, RR047A), according to the manufacturer’s protocol. Quantitative RT-PCR was performed with fluorescent SYBR Green on a CFX96 Touch System (Bio-Rad, USA). Human GAPDH was used to normalize input cDNA.

### Xenografts

Methods for xenograft implantation were described previously(Li et al, 2014). All animal protocols were approved by the Ethics Committee of the Third Military Medical University. Briefly, 2×10^6^ PC-9GR cells were injected subcutaneously into the back, next to the left forelimb of 6-week-old female BALB/c A-nu mice (Laboratory Animal Center of Third Military Medical University, Chongqing, China), which all developed tumors of ∼30 mm^3^ within 5 to 7 days. The mice were then randomly assigned to 4 groups (8 mice/group), and treated with CQ (100 mg/L), osimertinib (20 mg/L), combined CQ and osimertinib, and drinking water (vehicle). The tumor volume was calculated as (length×width^2^)/2, and measured twice a week. The animals were maintained in individual ventilated cages in compliance with institutional guidelines. The animals were monitored for 8 weeks until euthanasia. For immunohistochemistry assay, tumor bearing mice in each group were sacrificed after 4 weeks of drug administration, and tumors were harvested, fixed with 4% paraformaldehyde, and paraffin embedded.

### Immunohistochemistry

For immunohistochemistry, tumor sections were fixed with 4% paraformaldehyde overnight, paraffin embedded, sectioned, and stained with primary antibodies raised against Sox2 and LC3 (1:200). Sample scoring was performed by the H-score method that combines immunoreaction intensity and the percentage of tumor cells stained.

### Patient information, sample preparation and NGS sequencing

A retrospective investigation was performed by profiling plasma or tissue biopsies from 39 patients with acquired resistance to osimertinib using NGS. This study was approved by the Institutional Review Board of Daping Hospital, Army military medical university. All patients were provided informed consent to this study and gave permission to the entire study. Disease progression was confirmed in each patient according to Recist 1.1 criterion. Tissue DNA was extracted using QIAamp DNA FFPE tissue kit (Qiagen) according to manufacturer’s instructions. Circulating cell-free DNA was recovered from 4 to 5 ml of plasma using the QIAamp Circulating Nucleic Acid kit by Qiagen (Valencia, California, US). DNA shearing was performed using Covaris M220. End repair and A tailing was followed by adaptor ligation. The ligated fragments with size of 200-400 bp were selevted by beads (Agencourt AMPure XP Kit, Beckman Coulter, California, US), hybridized with probe baits, selected by magnetic beads and amplified by PCR. Indexed samples were sequenced on Nextseq500 sequencer (Illumina, Inc., USA) with pair-end reads.

### Statistical analysis

Statistical analysis was performed by GraphPad Prism 5 and the data were presented as mean ± S.E.M. The two-tailed Student’ s t test was used to compare multiple sets of data. P < 0.05 was considered to be statistically remarkable.

## Acknowledgments

We thank Dr. Raffaella Sordella from Cold Spring Harbor Laboratory for kindly providing the valuable cell lines used in the current study. We thank Prof. Xiang Xu and Prof. Xueqing Xu (both from Army Medical University) for their kind advice about design of the study. This work was supported by the Natural Science Foundation of China (81672284, 81472189, 81672287, 81702288 and 81702291) and a foundation for PLA Young Scientists (16QNP106).

